# A one-step CRISPR-based strategy for endogenous gene tagging in Drosophila melanogaster

**DOI:** 10.1101/2022.04.04.487042

**Authors:** Yangbo Xiao, Ye Yuan, Swathi Yadlapalli

**Affiliations:** Cell and Developmental Biology Department, University of Michigan, Ann Arbor, MI 48109, USA; Cellular and Molecular Biology Program, University of Michigan, Ann Arbor, MI 48109, USA

**Keywords:** CRISPR, endogenous gene tagging, single-leg non-lethal PCR

## Abstract

The ability to tag endogenous genes to create fluorescent fusion proteins has revolutionized the studies of protein subcellular localization and dynamics and regulation in live cells. Here, we report a precise, rapid, and one-step endogenous gene tagging system in *Drosophila melanogaster*, which allows us to screen for engineered lines without the visible eye marker. Specifically, we developed an efficient screening strategy for the identification of the homologous integration events by employing a PCR-based method to characterize individual flies using only a small segment of their middle leg. Here, we detail design and construction of gRNA plasmids and donor plasmid, and methods for screening, and confirmation of engineered lines. Together, these protocols make tagging any endogenous protein in Drosophila more efficient and faster, enabling studies of a wide range of cellular processes in live cells within intact organisms.

## INTRODUCTION

Fluorescent proteins have emerged as powerful tools for visualizing protein localization, dynamics, and protein-protein interactions in cells within living organisms^1^. Tagging endogenous genes with fluorescent proteins enable long-term live imaging of cellular processes at subcellular resolution. Furthermore, these endogenous gene-labeling techniques have several advantages compared to overexpression and immunostaining approaches. For example, overexpression of proteins can lead to mislocalization or aggregation artefacts, and static methods based on fixation can lead to the masking of epitopes that can prevent antibody binding. Till date, researchers have been able to create vast libraries of endogenous genes fused to fluorescent proteins for a few model organisms such as budding yeast, which can undergo efficient homologous recombination. The discovery of CRISPR-Cas9 based tools has only recently made it possible to efficiently perform genome editing, including tagging endogenous proteins, in any organism^2^.

CRISPR-Cas9 genome editing tools are currently being used to generate transgenic animals in which endogenous proteins are labeled with fluorescent proteins, enabling studies of cellular mechanisms in a wide range of organisms. Cas9 is an RNA-guided enzyme that cleaves double stranded DNA in a highly efficient and specific manner, which can then be repaired by nonhomologous end-joining (NHEJ) pathway leading to point mutations or deletions or homology-directed repair (HDR) pathway leading to integration of heterologous DNA into the chromosome. Previous methods for gene tagging in *Drosophila melanogaster* comprised of two-steps, where first a visible eye-color marker (Dsred) is used to identify a successful homology directed repair event and then the Dsred marker is removed by Cre/loxP recombination system^3^. Screening strategy typically involved examining hundreds of flies, where tens or hundreds of progeny of individual first-generation animals are molecularly screened to identify the homologous integration event. Here, we describe a much simpler procedure which allows non-lethal genotyping of individual animals by directly screening individual first-generation candidate CRISPR-flies and only establishing stocks from the appropriate individuals. This screening strategy will make screening for CRISPR-positive flies substantially faster, cheaper, and more efficient. Specifically, we developed a PCR-based protocol for genotyping single Drosophila by extracting DNA from a small segment of the middle leg. We found that using a small segment of the middle leg of a single, live fly for DNA purification and amplification does not have any effect on the fly’s viability, mobility, and reproductive ability.

Here, we present a detailed protocol for generating fluorescent protein tagged flies in which endogenous genes are labeled with fluorescent reporters. This technique uses CRISPR-Cas9 genome editing to fuse the fluorophore tag to the N- or C-terminal of an endogenous gene. Here, we describe the design and construction of gRNA plasmids and donor plasmid, one-step screening strategy to select CRISPR-positive flies from the first generation of injected flies, and confirmation of engineered lines. Specifically, we describe strategies to tag an endogenous core clock Drosophila gene, period, with a fluorophore at the C-terminal^4^. We also briefly describe the control experiments that need to be performed to confirm that the tagged protein is functional and live-imaging techniques to visualize the period protein in live clock neurons within intact Drosophila brains. The protocol described here is broadly applicable to also screen candidates for deletions and insertions of specific bits of DNA by CRISPR-Cas9 genome editing.

## PROTOCOL

### I) Design and construction of gene editing reagents

To conduct endogenous tagging, start with identifying different mRNA and protein isoforms of the gene. Typically, a fluorescent tag can be added to either N- or C-terminal of your targeted protein isoform(s), which will affect a) whether you can cover most of or specific protein isoforms, and b) whether the added fluorescent tag will affect normal function of the protein. Although it is desired to put the tag away from conserved regions of the protein, successful tagging can only be verified by a) existence of the tag sequence at the endogenous locus, and b) performing appropriate control experiments to test that the tagged protein is functional. To facilitate effective CRISPR editing, two sgRNAs need to be generated near the site of knock-in. Finally, to allow homologous recombination mediated editing, a donor vector containing homologous sequences to the target gene together with the gene tag need to be synthesized. Below we provide a step-by-step protocol for this procedure (Fig.1).

**Figure 1.**
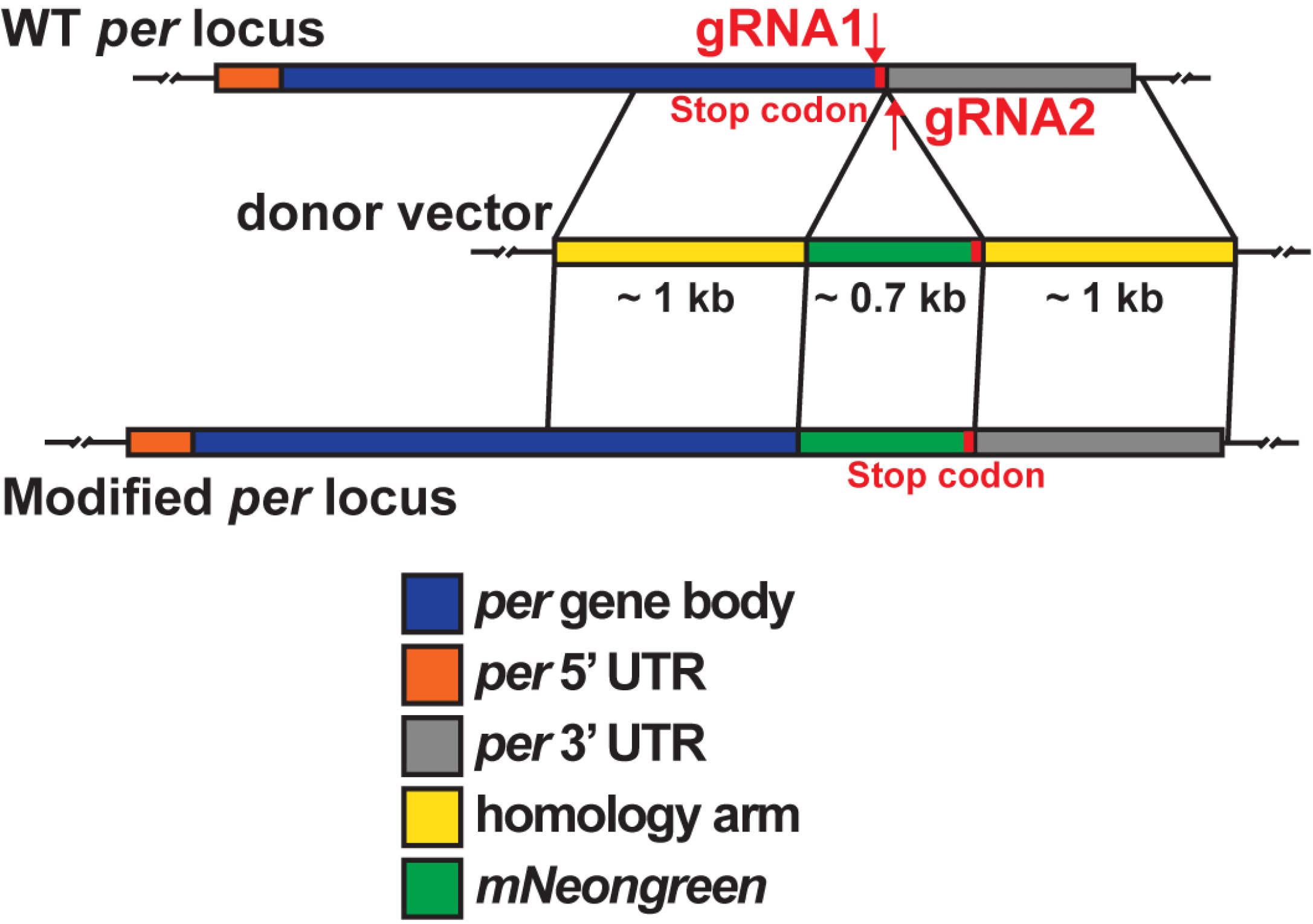
Schematic of CRISPR/Cas9 genome editing to generate *period-mNeonGreen* flies. Strategy for tagging the endogenous *period* gene with the mNeonGreen fluorescent tag. The two gRNAs are located on either side of the stop codon, one in the last exon and the other in the 3’UTR sequence. The donor vector contains two ∼1 kb homology arms and mNeonGreen tag with stop codon attached to it.

#### A) sgRNA design

An sgRNA is made up of a crRNA, which is a 20-nucleotide sequence corresponding to the region of the genome you wish to target, and a tracrRNA which scaffolds the sgRNA to Cas9. The tracrRNA is constant, while the crRNA is changed based on the region of the genome being targeted. Importantly, immediately adjacent to the 3′ end of the crRNA should be an “NGG” (protospacer adjacent motif (PAM)) sequence which allows for cleavage by Cas9 protein. Online tools examine the region of the genome you wish to target and look for 20 nucleotide sequences adjacent to NGG sites and then probe for relative off-target homology across the rest of the genome. Ideal crRNAs for tagging purposes will have minimal homology with the rest of the genome to decrease chances of off-target genome cleavage by Cas9. Here we will discuss how to identify optimal crRNA sequences using the online tool flycrispr target finder (https://flycrispr.org/target-finder/).

1. Using a web browser, go to https://flycrispr.org/target-finder/, click “Go to flyCRISPR Target Finder” button on the top.
2. Input ±100 bp of the site of knock-in. Select genome “Drosophila melanogaster (nos-Cas9 III, BDSC 78782)” if nos-Cas9 attP2 will be the injection background. Then choose: “CRISPR targets with 5’ G” option because crRNA sequence start with G would enhance the sgRNA expression efficiency. Click “Find CRISPR Targets” button. Nos-Ca9 allows Cas9 expression specifically in the germline.
3. Now you should see a list of potential crRNA sequences. Hit “Evaluate”. This step will look for potential off-targets using the previously selected genome and generally takes a few seconds.
4. Pick 2 crRNAs with “0 off targets”. Preferably, they should be as close to the knock-in site as possible to facilitate efficient editing. One crRNA should locate upstream of the knock-in site and the other crRNA downstream. crRNAs should not overlap or too close to each other (within 10 bp). Note: Ideal crRNAs are not always available depending on your gene of interest. We recommend prioritizing “0 off target” while other options are relatively flexible. The distance between crRNA and the knock-in site can be as far as 200 bp. crRNAs not starting with G are acceptable but will reduce the positive rate.

*IMPORTANT* Genomic sequence of the injection fly (y, sc, v genetic background) and the reference genome published by Flybase is different. To get accurate genomic sequences of the gene of interest, SNPs can be identified with the Find CRISPRs database (https://flyrnai.org/crispr3/web).

#### B) Fluorescent tag selection

5. There are many suitable fluorescent tags with strengths and weaknesses^5^. When selecting a fluorescent tag to label your protein of interest, consider the following:
  a. Excitation/emission wavelengths of your tag – Choose a tag that you can easily image with equipment accessible to your lab and that is compatible with other assays you plan to perform once you have generated the knock-in flies.
  b. Stability and intensity of the fluorescent tag – Choose a fluorescent tag that will not quickly photobleach and that is relatively bright because the amount of endogenous protein is usually much less than overexpression assay^5^.
  c. Toxicity of the fluorescent tag – Some fluorescent tags, such as tdTomato and DsRed are known to aggregate, and thus using these fluorescent tags as your tag may be undesirable (Bell et al., 2019). Choose a fluorescent tag with minimal toxicity. Note: Red fluorescent proteins such as mScarlet-I tend to be stable even in acidic environment (e.g., in lysosome)^6^. Therefore, if the target protein is mainly cytoplasmic, choosing other available fluorophores would be preferred. Note: Other small peptide tags (e.g., FLAG, V5) can be attached to the fluorescent tag bridged by a linker to enable biochemistry assays such as Western Blot or Co-Immunoprecipitation.

#### C) Linker design

6. It is often desirable to have a linker peptide that connects your fluorescent tag to your protein of interest. This increases the chances that the fluorescent tag and endogenous protein of interest will properly fold and wildtype function and localization are retained. Linker peptide sequences are normally 2–20 amino acids long and typically consist of amino acids such as glycine and serine to provide flexibility^7^. In our design, we found that double glycine linker is sufficient to preserve the activity of our target protein (Period).

#### D) Donor vector design

7. The donor vector contains the tag you wish to place into the genome flanked by sequences homologous to the gene of interest. To design the donor vector, arrange the linker peptide and fluorescent tag according to whether they will be placed at the amino or carboxyl terminus of your protein of interest and then add homologous sequences of DNA on both sides of the knock-in site. We recommend using homology arms that are at least 800 nucleotides long on each side of the knock-in position. Note: If your fluorescent tag is placed at the amino terminus of your protein of interest, be sure to remove the stop codon from the fluorescent tag so that translation does not stop prematurely. Remove the start codon in the target gene coding sequence so that translation could only start at the fluorescent tag site. Conversely, if your fluorescent tag is placed on the carboxyl terminus, place your fluorescent tag upstream of the stop codon of the endogenous gene so that translation does not stop prior to your fluorescent tag. In this situation, be sure to place the stop codon in the repair template to make sure translation ends correctly and in frame. Remove the first methionine from the fluorescent tag sequence for carboxyl terminus tagging.

#### E) Synthesize the sgRNA expression plasmids and the donor plasmid

8. For generation of gRNA plasmids, we followed U6-gRNA cloning protocol to generate gRNA plasmids (https://flycrispr.org/protocols/grna/). Specifically, we used the alternative restriction enzyme cloning strategy (strategy C in the reference). From IDT DNA, we ordered 2 single-strand DNA oligos for each gRNA. Then, we annealed oligos and phosphorylated the annealing product. Then we followed the ligation protocol. After ligation, we did transformation (NEB C2987H) and checked the transformants by Sanger sequencing. For generation of the donor plasmid, individual gBlock sequences containing fragments of the donor sequence can be ordered. The full donor sequence can be cloned via Golden Gate Assembly (citation needed) or other common cloning methods. We use pBlueScriptII SK(-) as donor plasmid backbone. Alternately, the whole donor plasmid can be ordered from a commercial vendor. It is important that the PAM sequences in the donor plasmid are either mutated such that the amino acid sequence of the protein is not altered or removed in the case of 3’UTR. If the PAM sequence is not altered in the donor plasmid, the sequence can be repeatedly cut by Cas9, which can lead to inappropriate sequences inserted into the genome.

### II) Injection and Screening using single leg PCR

The injection requires Midi-prep sample (QIAGEN). The concentration for gRNA should be over 0.5μg/μl and over 1.0μg/μl for donor. Injection can be done through commercial vendors or can be done in-house. The company will ship back the injected embryos in ∼2 weeks. The flies from company (G0) only have the CRISPR-edited allele in their germline cells. Therefore, we need to cross them with balancer flies first. Male G0 flies need to be crossed with balancer virgins individually while 10∼15 female G0 flies can be grouped in one vial and crossed with balancer male flies. We developed a single-leg PCR protocol which enables us to genotype an individual fly from a small segment of the single middle leg and use that fly for crossing with balancer lines, if that fly possesses the desired CRISPR allele. This will allow us to efficiently screen the transformants in the first round itself and only those flies which are positive in that first round will be crossed to make stable lines. Here we show an example of a gene on the X chromosome. Editing gene on second or third chromosome would require using corresponding balancer lines.

9. From the injected flies, pick female virgins and males – we can usually collect about 30 of each – We call them G0.
10. G0 male crosses – Let us call them ‘A’ crosses
  a. Set up crosses with individual G0 male and 2 or 3 FM7/L female virgins.
  b. You need to set up as many crosses as the number of G0 males. We do this to increase chances that each male has offspring. (If you group 4 or 5 males in 1 cross, females might only mate with one of the males)
11. G0 female crosses – Let us call them ‘B’ crosses. You can group about 10 G0 female virgins and 4-5 FM7/Y (X chromosome balancer) male flies. All the females will mate and produce offspring even if you group them in a vial. We usually set up about 2-3 ‘B’ crosses.
12. You need to now start the screening process with the F1 progeny from ‘A’ and ‘B’ crosses. – Collect female F1 virgins from ‘A’ and ‘B’ crosses. Collect F1 males from ‘B’ crosses.
13. From the ‘A’ crosses, you can only use F1 females. These F1 females will potentially have your CRISPR-modified gene on one X chromosome and another X chromosome will be FM7-positive. F1 males from ‘A’ crosses are not useful as they get their X chromosome from FM7/L moms.
14. From the ‘B’ crosses, you can use both F1 females and F1 males. These F1 females can potentially carry your CRISPR-modified gene on their X chromosome.
15. We usually start screening F1 males and then move to F1 females. We can screen about 50-60 flies every day. We stop screening when we find ∼50 CRISPR-positive flies.
16. Single leg PCR protocol:
  a. Primer design: we recommend one primer in the 5’ homology arm of target gene and another primer in the 3’ homology arm of the target gene for initial screening because the CRISPR-positive flies will generate 2 different bands in agarose gel (heterozygous) while the CRISPR-negative flies and the control flies will also give a lower band which indicate the screening process is correct.
  b. For each fly to be screened prepare the lysis buffer in 96-well PCR plate:

  c. Prepare glass capillary tubes (with sucrose-agar food, 2% agar and 4% sucrose – same as DAM behavior tubes) and place them in a 96-well deep-well plate (referred to as “fly hotel”) (Fig.2).
  d. Under the microscope, carefully cut a small segment of the middle leg of the fly by forceps and place it into the lysis buffer in 96-well plate. Make sure it is soaked in the buffer (underneath the liquid surface). Cutting part of the middle leg doesn’t affect the fly’s mobility or fertility.
  e. Take the fly and place it into the corresponding well of “fly hotel”. Cap the behavior tube with yarn (Fig. 2).

| Chemical Name | Stock Conc. | Volume |
| --- | --- | --- |
| Tris HCl (pH8.0) | 1 M | 0.1 μL |
| NaCl | 2.5 M | 0.1 μL |
| EDTA (pH8.0) | 0.1 M | 0.1 μL |
| Proteinase K | N/A | 0.1 μL |
| Nuclease free water | N/A | To 10 μL |

**Figure 2.**
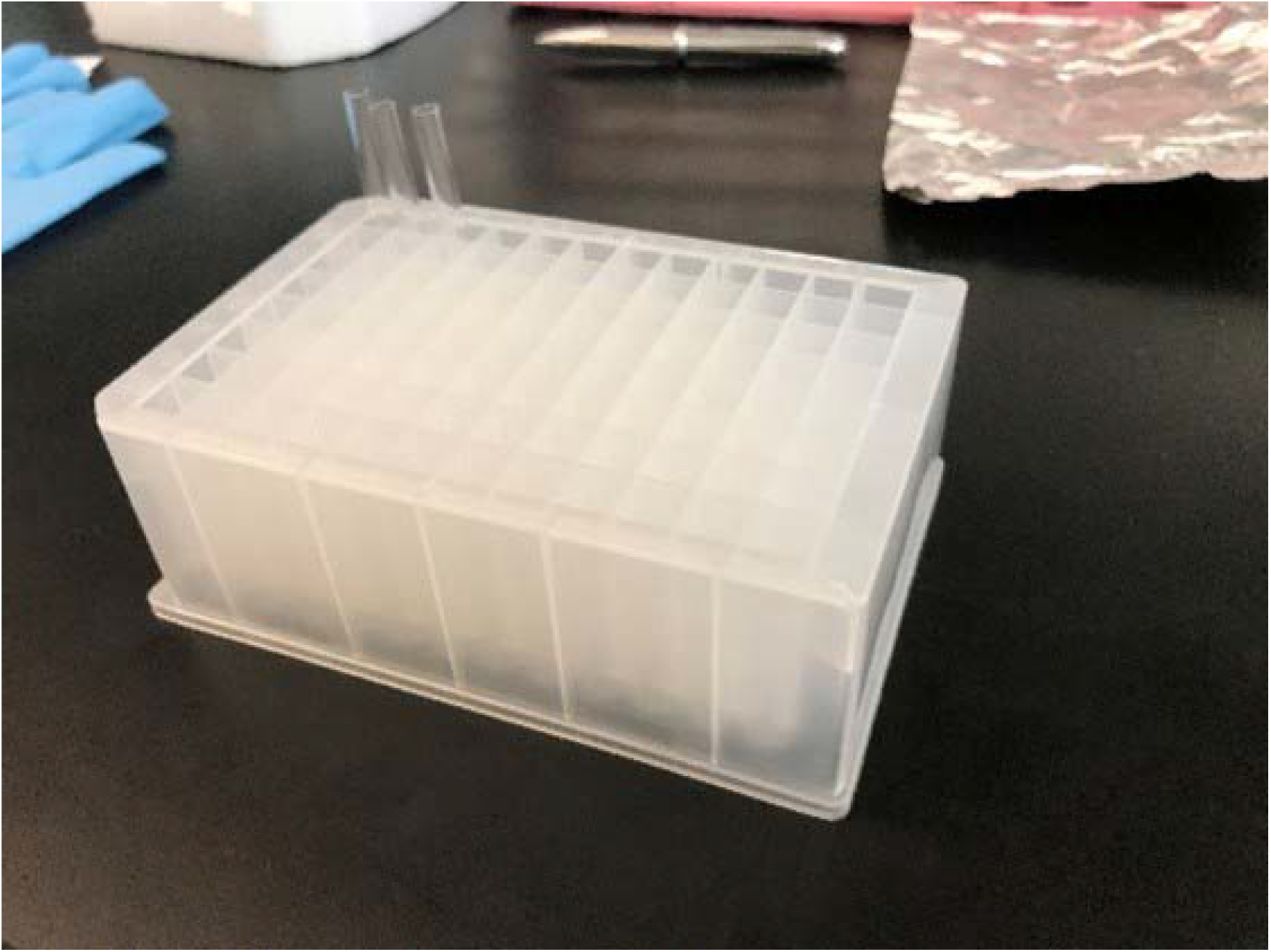
Picture of the ‘Fly Hotel’. Glass tubes are filled with fly food and sealed at one end. Single flies are loaded into individual tubes. After the DNA extracted from the middle led of individual flies is tested, the appropriate flies are taken from the fly hotel and the lines are established.
  f. Remember to dissect 2 flies for negative control and 2 for positive control. If no flies can serve as positive control, the donor plasmid can be diluted and used instead.
  g. When leg dissection is done, place fly hotel in 25-degree incubator. Flies can survive overnight, but it is better to finish the following steps and cross them within one day.
  h. Run the following program in a thermocycler:

  i. Prepare the PCR reaction for each sample as following:

| Temperature | Time |
| --- | --- |
| 37°C | 90 min. |
| 95°C | 5 min. |
| 4°C | hold |

  j. Run the following program:

i. Prepare the PCR reaction for each sample as following:
|  |  |
| --- | --- |
| 2X PCR master mix with dye | 10μL |
| 10μM forward primer | 1μL |
| 10μM reverse primer | 1μL |
| Leg lysate | 3μL |
| Nuclease-free water | to 20μL |

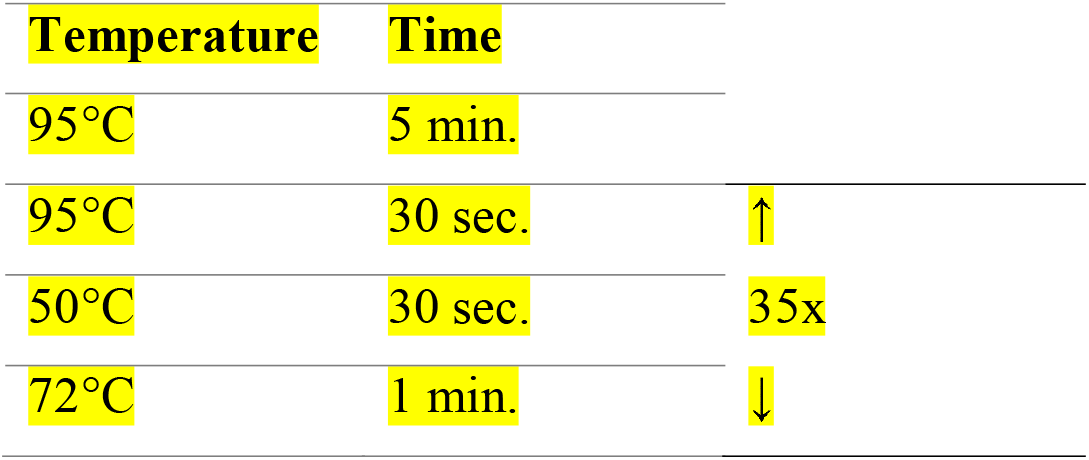
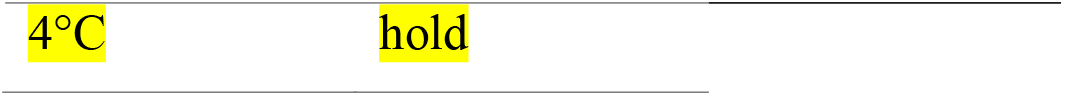
  k. Run the PCR products in agarose gel.
17. Pick the positive flies from “fly hotel” and cross them with balancer flies to generate F2 flies.
18. When F2 flies eclose, do sibling crosses to generate stable stocks.
19. *optional* Single leg PCR can be applied to F2 flies with different sets of primers (e.g. one in the 5’ homology arm another in ∼100bp downstream of 3’ homology arm region) to check the location of the insertion in the endogenous gene locus to ensure that the whole donor plasmid is not inserted in some random location in the genome.
20. Pick homozygotes from F3 flies. Extract genomic DNA and do sanger sequencing to confirm the sequence of the fly of target region.
21. If desired, back cross the confirmed fly line with WT flies to remove background mutation. Every generation, flies can be screened by single leg PCR.
22. Appropriate control experiments need to be performed to check that the tagged protein is functional (More details provided in Representative Results section).

j. Run the following program:
| Temperature | Time |  |
| --- | --- | --- |
| 95°C | 5 min. |  |
| 95°C | 30 sec. | ↑ |
| 50°C | 30 sec. | 35x |
| 72°C | 1 min. | ↓ |

|  |  |
| --- | --- |
| 4°C | hold |

### III) Live imaging

After successful generation of transgenic flies in which endogenous protein is fluorescently tagged, we can perform live imaging to study the subcellular localization and dynamics of the fluorescently tagged protein in cells within live tissues^4^. As an example, here we describe the live imaging protocol for tagged clock proteins in adult Drosophila brains.

Flies were entrained to Light-Dark (LD) cycles with lights on for 12 hours and off for 12 hours for 5-7 days, and then released into complete darkness (DD). All flies used for live imaging experiments were placed in food individual vials (4 females and 4 males) and entrained for 5-7 days in incubators. For imaging of clock proteins in live clock neurons, 3-4 adult brains were dissected in chilled Schneider’s *Drosophila* medium in less than 5 minutes. A punched double-sided tape was used as a spacer on the slides to prevent flattening of the brains. The brains were overlaid with a small amount of Prolong Glass Antifade mounting medium and covered using a coverslip. We acquired individual z-stack images of clock neurons using the Zeiss LSM800 Airyscan laser scanning confocal microscope. We have acquired our images using a 63x Plan-Apochromat Oil (N.A. 1.4) objective and 405, 488, and 561 nm laser lines. We collected Z-stack (each Z-slice is about 250 nm) or time-lapse image series of individual clock neurons and images were analyzed using Zeiss ZEN software and ImageJ.

## REPRESENTATIVE RESULTS

This method can be used to tag endogenous genes on any of the chromosomes in *Drosophila melanogaster*. In our experiments, we have successfully used these strategies to tag genes all the chromosomes (X, II and III chromosome). The positive rate varies depending on gRNA selection, characteristics of the gene, and efficiency of injection. Here we provide representative results of the screening strategy for endogenous gene tagging of *period* gene with mNeonGreen tag^8^. *period* is a gene located on X chromosome of *D. melanogaster*, which is a key component of circadian clock.

*Drosophila* has a simple and stereotyped clock network consisting of 150 clock neurons, which express clock proteins^9^. We tagged Period protein with green fluorescent protein mNeonGreen at C’ terminal bridged by a double glycine linker. After plasmid injection, we crossed the flies to FM7/L or FM7/Y and collected F1 flies. We screened 126 F1 flies using PCR primers (see material table) and 47 of them are positive. The positive rate is 37.3%. We did not observe any significant positive rate difference due to gender in the screening process. After confirming that the sequence of the tagged gene is correct, we performed control experiments (behavior, qPCR to determine mRNA oscillations) to check that the tagged protein (PERIOD-mNeonGreen) is functional by checking that the protein and mRNA levels are oscillating similar to the endogenous gene in wild-type flies and that the behavior of the flies is rhythmic^4^. We then crossed these PERIOD-mNeonGreen flies to *CLOCK*^*856*^*-GAL4;UAS-CD4-tdTomato* flies (*CLOCK*^*856*^ is a pan-clock neuron marker) to image PERIOD subcellular localization and dynamics in individual clock neurons in flies entrained to light-dark cycles^4^. Strikingly, we observed that PERIOD protein is concentrated in distinct, dynamic foci in the nucleus during the circadian repression phase. Representative images for PERIOD protein in live clock neurons in Drosophila brains are shown in Fig. 3.

**Figure 3.**
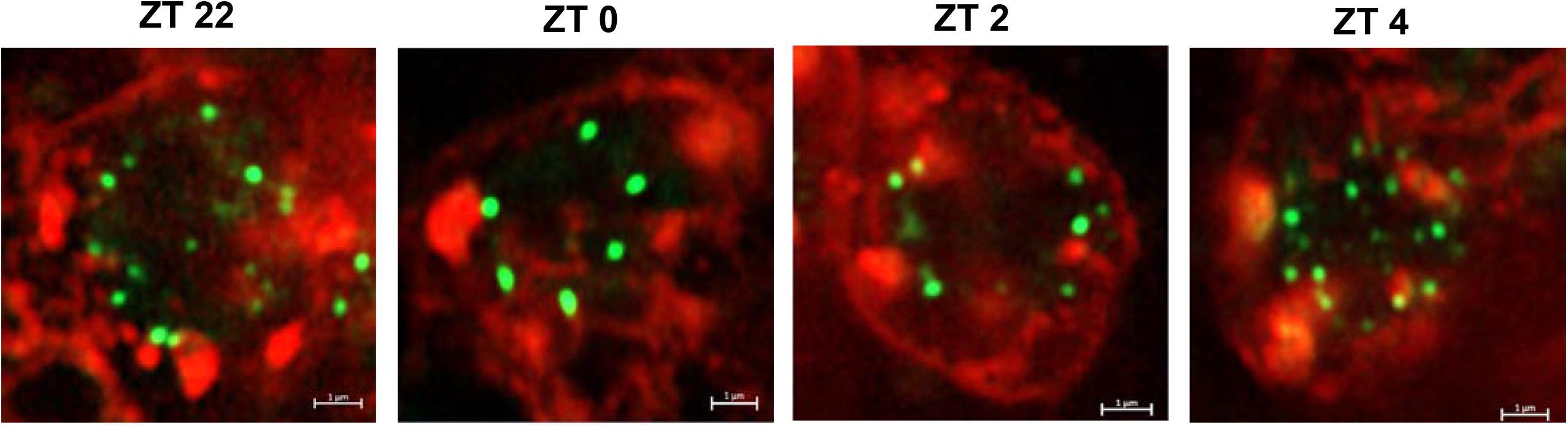
Representative images of PERIOD-mNeonGreen protein in clock neurons. Images from *period-mNeonGreen;Clk-GAL4,UAS-CD4-tdTomato* flies entrained to Light-Dark cycles where lights are turned on at ZT0 and turned off at ZT12 (‘ZT’ refers to ‘Zeitgeber Time’). PERIOD protein is organized into discrete perinuclear foci during the circadian repression phase (ZT22, ZT0, ZT2, ZT4). Representative images of PERIOD foci (green) in a group of clock neurons, sLNvs (cell membrane labeled with tdTomato and shown in red), over the circadian repression phase. Scale bars, 1 μm.

## DISCUSSION

Our results demonstrate a streamlined, simple one-step CRISPR-based strategy for tagging endogenous genes in Drosophila. This method provides significant advantage over the current screening methods which use two steps of screening, where first step relies on using a visible marker such as eye color to identify the appropriate genotype and the second step involves removal of the eye color marker^3^. Here, we describe a non-lethal PCR on DNA extracted from a small segment of the middle leg of an individual fly allowing genotyping at least one generation earlier, which can replace current molecular methods with a significant saving in time and cost. This method allows screening of candidate animals one by one or in small batches (up to 60-70 can be screened at a time without any issues) until the appropriate genotype is isolated. The number of animals screened before appropriate stocks are isolated varies from experiment to experiment, depending on many factors including efficiency of sgRNAs selected, efficiency of injection, off-target insertions which can affect viability. It is also possible the addition of a tag to the N- or C-terminal of a gene can adversely affect the function of the gene, which can potentially affect the viability of the flies. If the protein structure information is available, it can be used to determine whether N- or C-terminal tagging will better preserve the protein’s function and localization of the native protein. It is always recommended to perform control experiments, including testing whether the tagged protein shows similar localization pattern as the wildtype protein by antibody staining methods. In practice, we typically found 10-30% success rate in screening for various endogenous tagged genes on all chromosomes (X, II, III). The same screening strategy can be used to screen to other kinds of CRISPR-based genetic insertions or deletions. Other published methods have used single wings from live flies to screen for chromosomal recombination and transposable element mobilization events^10^.

In conclusion, our one-step screening strategy makes it much easier to generate CRISPR-positive flies and opens many possibilities to study the subcellular localization and dynamics of endogenous genes in their native context.

## ACKNOWLEDGEMENTS

We thank George Watase and Josie Clowney for discussions during the initial stages of the protocol development. The work was supported by funds from the NIH (grant no. R35GM133737 to S.Y.), Alfred P. Sloan Fellowship (to S. Y.) and McKnight Scholar Award (to S. Y.).

## MATERIALS

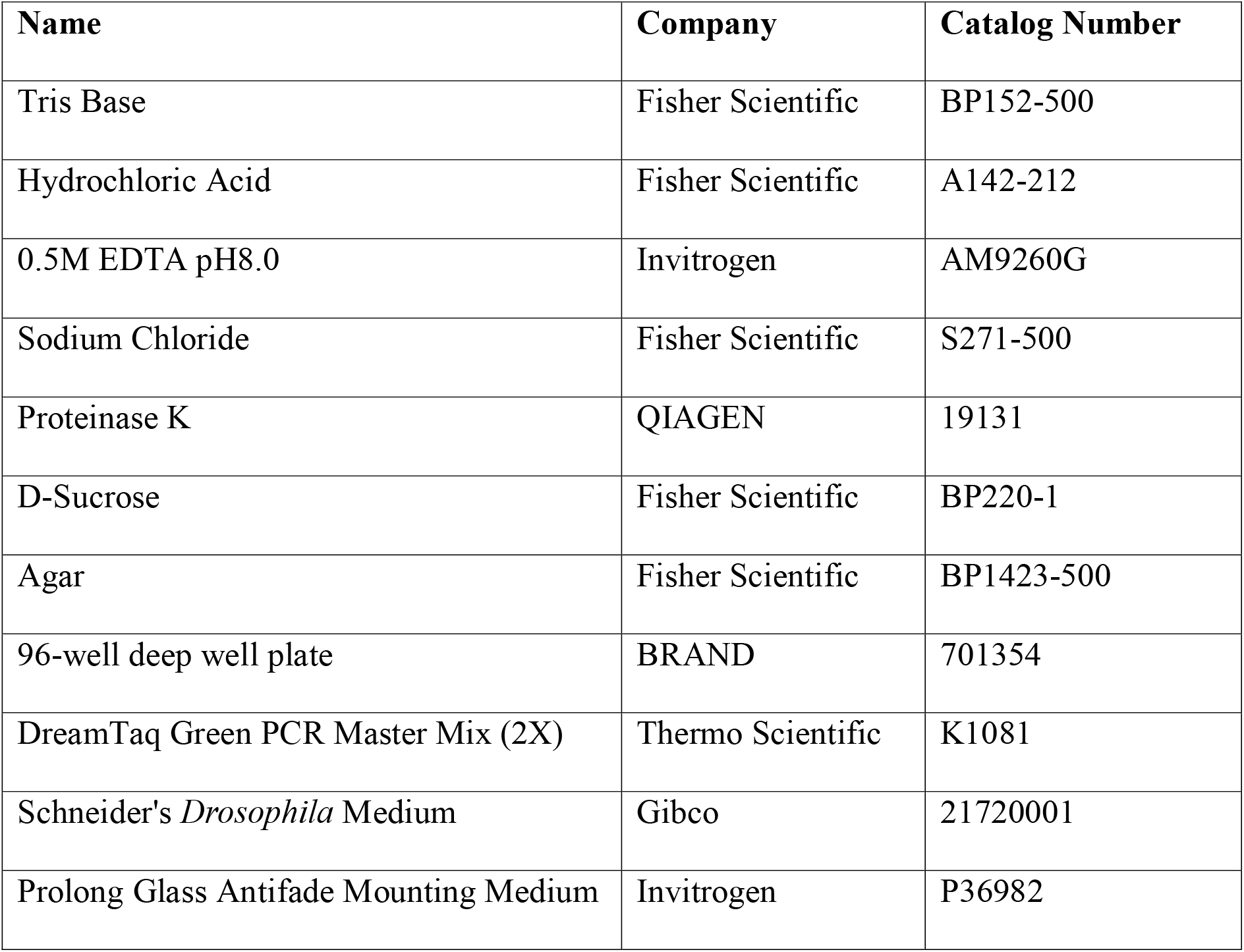
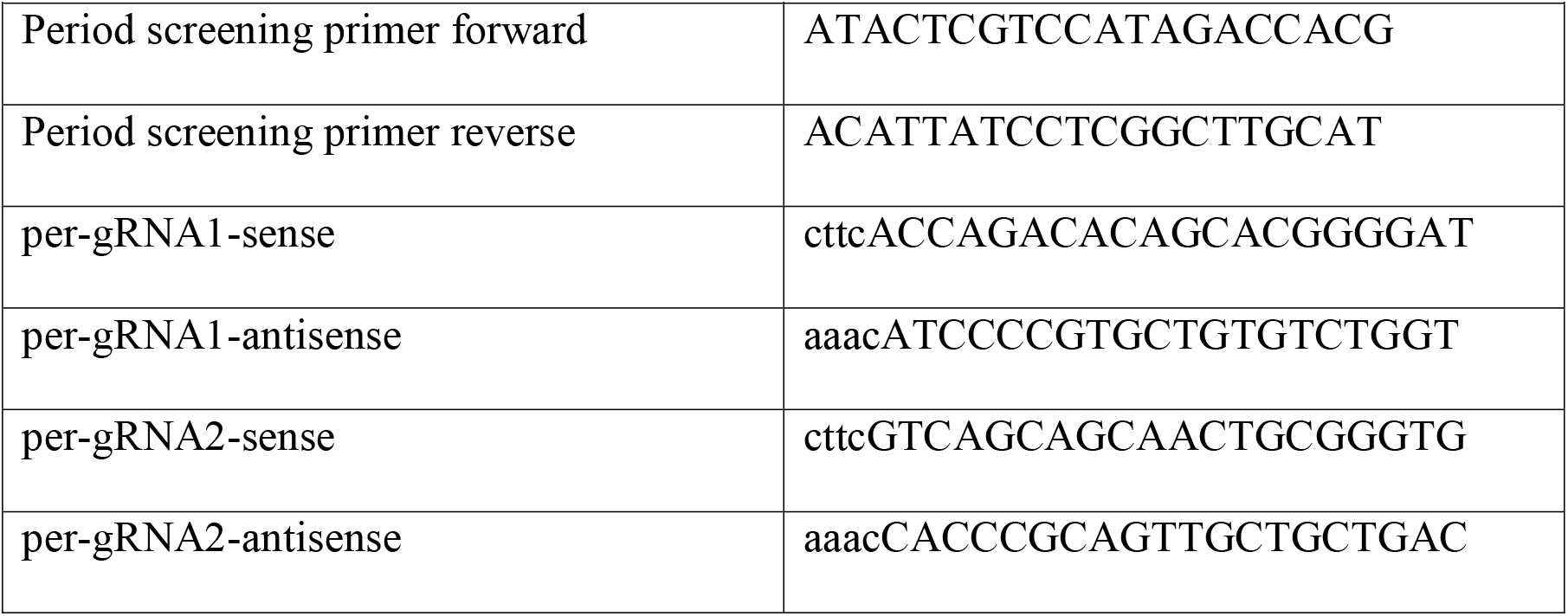

